# Multilevel-consistency of social behaviour in a cockroach

**DOI:** 10.1101/2025.08.29.673046

**Authors:** Callum J McLean, Sanskar Amgain, Binod Bhattarai, David N Fisher

## Abstract

Group-living animals may show consistency in both individual behaviour and group-level behaviour. Further, while individuals show consistent differences in their responses to environmental change (plasticity), it is less clear if groups differ in plasticity. Here we exposed groups of cockroaches (*Blaptica dubia*) to three humidities and quantified their social networks. We repeated our design across multiple groups with individuals reassigned to different groups and used machine vision analysis of images to extract 152,800 records of associations across 16 different groups. 116 unique individuals contributed >4,500 individual social network measures and 192 measures of entire networks. Lower humidity led to individuals that had slightly better connectedness to the whole network, but otherwise did not impact individual or group-level phenotypes. Individuals showed some consistency in mean behaviours (12-39%) but scant consistency in plasticity (0-9%). In contrast, groups showed less consistency in mean behaviours than individuals (0-26%) but showed relatively higher among-group differences in plasticity compared to individuals (0-22%). Patterns of group-level variance in individual traits matched patterns of variance in characteristics of entire networks, suggesting these two approaches for quantifying group phenotypes are compatible. While consistency in individual mean behaviour is the norm across taxa, consistency in group-level plasticity is less well understood and may be an emergent phenomenon of collective dynamics that deserves further investigation. Using automated marker recognition techniques such as machine vision allows us to collect the large datasets necessary to simultaneously test hypotheses at the individual and group level and so can be more widely adopted.

## Introduction

The behaviour of animals is typically repeatable i.e., traits such as boldness and sociability are consistently different among individuals across contexts and over time (Bell et al., 2009). Alongside these differences in mean behaviour, animals also may differ in how they respond to the environmental factors (Dingemanse et al., 2010; Nussey et al., 2007). Some individuals may increase a trait as temperatures warm, group size changes, or as they age, while other individuals may show no change or a decrease. This individual variation in plasticity has important consequences for ecological and evolutionary processes (Dingemanse & Wolf, 2013), as diversity in responses to changes in external conditions may buffer populations against perturbations (Maldonado-Chaparro et al., 2017), for instance by increasing the probability that some individuals in the group achieve optimal responses (the “portfolio” effect; Schindler et al., 2015). Additionally, individual variation in plasticity sets the upper limit for the heritability of plasticity, which is key for evolutionary change in flexibility (Pigliucci, 2005).

Behavioural consistency is not limited to individuals. Animals that live in groups may show collective behaviour; moving and acting as a cohesive unit (Sumpter, 2006), to enhance thermoregulation (Joos et al., 1988), improve hunting success (although this depends on how hard prey are to capture; MacNulty et al., 2012, 2014), reduce requirements for individual vigilance (Beauchamp, 2008), and improve decision making (Sumpter & Pratt, 2009). Collective behaviour is therefore considered an important phenomenon in behavioural ecology and even in human psychology (Hofmann & Jones, 2005).

Recent work has shown that groups differ in their mean behaviours much like individuals (Bengston & Jandt, 2014; Ioannou & Laskowski, 2023). For example, shoals of sticklebacks (*Gasterosteus aculeatus*) change their behaviour across contexts, but still show differences among groups in speed, alignment, cohesion, and consistency of the identity of the lead fish (Jolles et al., 2018). This among-group consistency in behaviour is subject to similar ontogenetic and ultimate explorations as personality at the individual level (Bengston & Jandt, 2014; Ioannou & Laskowski, 2023). It is also therefore logical to explore whether groups show consistent differences in their responses to environmental stimuli. Such responses could spring from the plasticity of individuals within the group or small initial differences could be amplified by positive feedback loops (Bengston & Jandt, 2014; for example see: MacGregor & Ioannou, 2021). Group-level variation in plasticity could provide another buffer for populations against perturbations (Gordon, 2013), and be a target for multilevel selection to act upon.

Group level behaviour and plasticity can be assessed at multiple levels, individuals in a group can have their behaviour averaged to give a measure of the collective phenotype, such as average aggression (Eldakar et al., 2010). Additionally, some behaviour of the group itself can be measured, such as degree of cohesion (Butail et al., 2014), which is not measurable at the individual level. These two approaches to measuring group-level phenotypes are however rarely conducted together and so are not directly compared for whether they show similar patterns of variation, for instance showing equivalent responses to stimuli or similar magnitudes of among- and within-group change. With interest in quantifying collective behaviour high it is key we understand if summarising across individual-level phenotypes and measuring group level phenotypes are comparable.

Testing for group-level variation in plasticity requires a large amount of data. Groups need to be sufficiently large to allow collective dynamics to emerge, there needs to be sufficient replication of groups to give the power to reliable detect among-group differences, these groups need each to be tested across multiple environments, and ideally individuals need to be mixed periodically between groups to separate the influence of individual personalities, especially highly influential “keystone individuals” (Modlmeier et al., 2014), from group-level personalities. Such challenging data requirements can be addressed through recent developments in automated marker recognition technologies that making gathering large amounts of data quickly across many individuals feasible (e.g. Dankert et al., 2009). These can be paired with manipulations of the environment or observations across natural environmental variation, and hierarchical mixed-effects models to quantify individual and group-level response to change, to measure consistency in mean group behaviour and consistency in group response to external stimuli.

Here we used a machine vision algorithm to quantify the social behaviour of 16 groups of the cockroach *Blaptica dubia*, with each group exposed to a gradient of decreasing relative humidities. Cockroaches naturally form aggregations that show collective behaviour (Freeberg & Fiset, 2023; Lihoreau et al., 2012) and are sensitive to humidity due to the risk of water loss (Dambach & Goehlen, 1999; Gunn, 1933; Yoder & Grojean, 1997) and so are an ideal study species for our goals. Along with individual variation in social behaviour (quantified as social network traits), we also quantified group-level variation as network level metrics (Croft et al., 2008; Whitehead, 2008). We assayed groups in blocks of four and mixed individuals between groups after each exposure gradient, allowing us to separate individual behaviour and group-level behaviour. We could therefore answer the following questions:

1. What are the population level responses of individuals and groups to decreases in humidity? We predicted that individuals would cluster more at lower humidities (as seen in other cockroach species; Dambach & Goehlen, 1999), leading to more dense and more tightly clustered social networks.
2. Do individuals and groups differ in their mean behaviour and in the plasticity of their behaviour? We predicted that individuals and groups would differ in mean behaviour (e.g. Jolles et al., 2017, 2018), and individuals would differ in their plasticity, but we did not make a prediction for group-level plasticity.
3. Do network-level traits show the same patterns in flexibility and consistency as corresponding group-level variation in individual-level traits? We predicted that, as both phenotypes are based on the same network of interactions, the patterns would be equivalent at different levels.

## Methods

### Trial Structure and study organisms

We assayed social behaviour in groups of 24 adult female *B. dubia* (Serville, 1839). We performed a total of four trials, each consisting of four groups. Trials took place over a three-week period, where we decreased humidity in a stepwise fashion over the trial, starting at 50% relative humidity (RH) in week one, 40% in week two, and 30% in week three. We kept the temperature at a constant 28°C over the trial and set the lights to a 12-12 photocycle. Between each trial there was a period of one week where we maintained the groups in the same conditions as the stock population (28°C and 50% RH; for further details on maintenance of the stock population see Fisher, 2023).

The groups for the first trial were comprised of cockroaches which we selected haphazardly from four stock boxes; we used a different stock box for each of the four first groups so that cockroaches within a group were initially equally familiar with each other. We assigned each individual with one of 24 unique letter tags (one of “2”, “4”, “5”, “6”, “7”, “a”, “b”, “d”, “e”, “f”, “g”, “h”, “i”, “J”, “k”, “L”, “N”, “Q”, “R”, “s”, “t”, “v”, “x”, or “y”, see below). We used the same 24 tags in each group. If an individual died during the trial it was replaced with another individual from the stock population with the same tag, keeping the group size at 24 (this occurred 73 times over the entire 15 weeks the experiments ran). We did however identify these as distinct individuals for the analysis. After the first trial we created new groupings by randomly assigning cockroaches with each tag to one of the four groups in the next trial, so that there was only one of each unique tag per group.

### Tagging

To ensure individuals could be correctly identified by the machine vision algorithm we attached two of the same tags made of waterproof paper measuring 75 × 75 mm each to the prothorax and abdomen, increasing the change at least one tag would be visible when a cockroach was in an aggregation. Each tag displayed a single letter (lower or upper case) or number to denote the ID of the individual (see list above). We selected a set of relatively distinct 24 tags across numbers, lower case letters, and upper-case letters to aid the machine vision algorithm and reduce the number of incorrect identifications. We only replaced tags when then fell off or became unreadable; tags remained in place between trials. We also gave all subjects an identifying marker using paint dots (Edding 780 extra fine paint marker) for researchers to identify individuals in case where both papers tags fell off. These paint marks were renewed at the start of every trial.

The impact of the tags used in this study on the movement and exploration of individual adult female *B. dubia* cockroaches was previously evaluated in McLean and Fisher (2024). That previous study found that the average speed of movement, the number of discrete walks, and the number of unique spatial zones visited over a 20-minute trial period was higher in tagged than untagged individuals, indicating that these tags have a positive effect on cockroach movement. This could have exacerbated any effects of changing humidity on cockroach movement, aggregation formation, and therefore social networks, but since the effect of humidity was modest (see Results), we do not think this is likely to be a problem.

### Experimental set-up

We carried out trials in a KBF P 240 incubator (BINDER GmbH), within which there were four shelves with lights above shelves one and three. On a shelf we assayed a single group in a day, and we rotated groups by one shelf for each day of recording. Groups were assayed on four consecutive days in a week (Tuesday-Friday) at a single humidity and then kept in the incubator at the same humidity for three days (Saturday-Monday) before the humidity was adjusted downwards and the groups assayed for social behaviour. After the trial at 30% RH was complete, we returned groups to the room with the stock population (see above). We carried out trials in clear polypropylene boxes (456 × 356 × 120 mm [l × w × h]). In each corner of the arena, we created a ‘hide’ measuring 150 × 100 mm by covering the exterior walls and lid of the box with opaque tape and so providing an area of shade. We also cut a 150 × 100 mm hole in the centre of the lid of the box and replaced it with a mesh covering to ensure sufficient ventilation. During the trials we supplied each group with 10g of carrot and 10g of hydrated polyacrylamide gel (Bug Gel, ProRep) per day Monday-Friday, placed in the centre of the boxes away from the hides. On Friday, 30g of carrot and 30g of hydrated gel were added to the arena to maintain hydration over the weekend; no carrot or gel was added on Saturday or Sunday.

To record the interactions between individuals we placed a V50 × camera (AKASO Tech) directly above each of the ‘hides’, cutting a hole in the box lid for the camera lens. We set the cameras to take photographs at one-minute intervals, capturing the identities of the individuals in the aggregations that had formed under each hide (Fig. 1). We recorded images on the first four days of each week of the trial for a period of four hours; we typically started filming between hours 6-7 of the daylight portion of the photocycle and ended between hours 10-11. This resulted in a maximum of 240 images per camera per day. However, due to battery depletion or other faults this number was not always reached, thus we always took the first 200 images of the trial (800 images per group per day as there were four cameras per group), equating to a recording period of 3 hours 20 minutes. We lost the data from one camera for one week (800 images) due to a corrupted memory card, but since this would be spread evenly across all groups (as the same cameras were used on a shelf while groups moved among the shelves) we do not correct for it. Our final set of images was therefore 200 images per camera (4), per group (4), per day (4), per humidity (3), per trial (4), minus the missing 800 images = 152,800.

**Figure 1.**
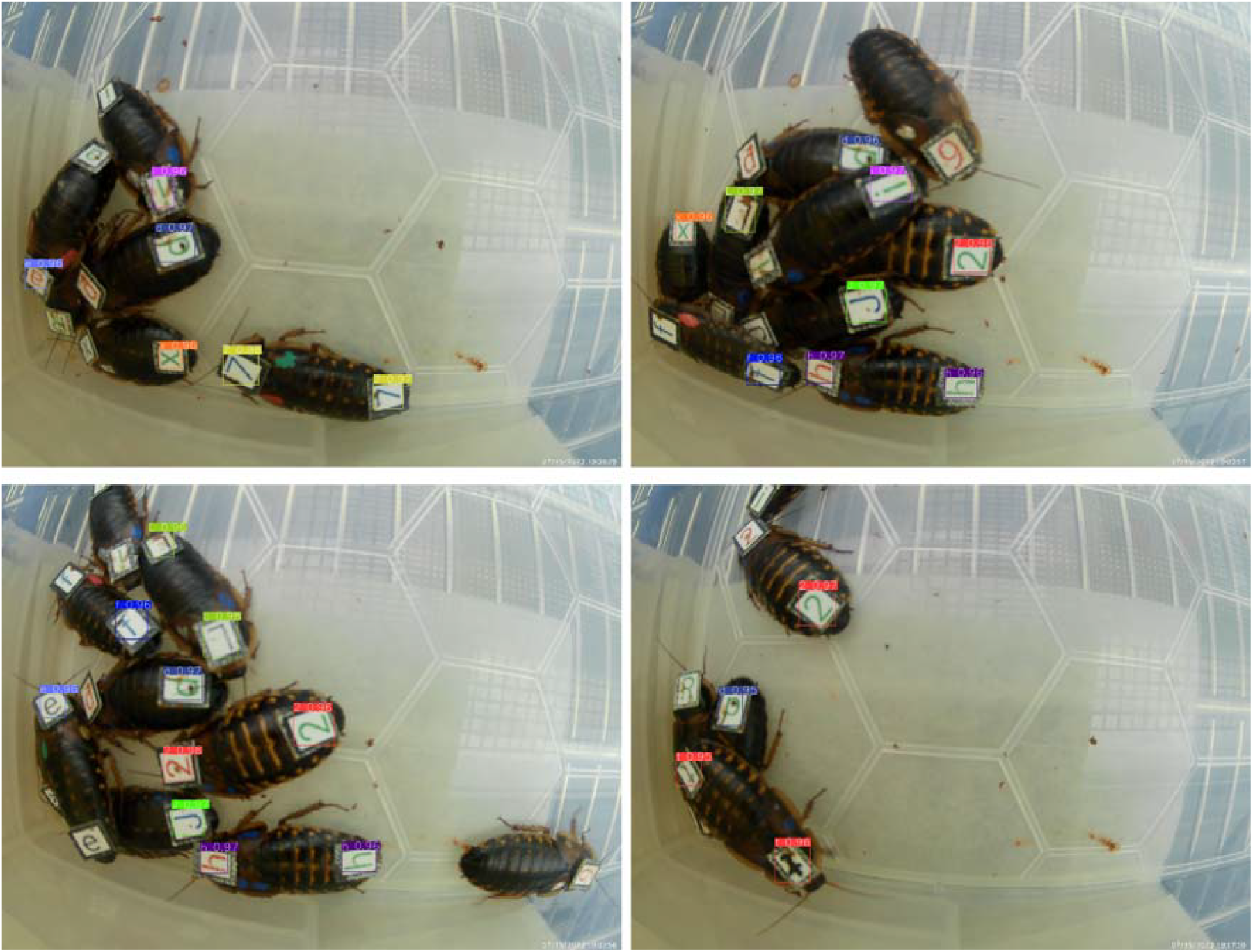
Four sample images of cockroach aggregations from the test dataset, with bounding boxes displaying the YOLOv5x predictions. The model’s confidence in each detection is displayed next to each bounding box.

### Tag Detection

To identify tagged cockroaches in images captured during the trials, we developed a specialized object detection algorithm based on the YOLOv5 extra-large (YOLOv5x) model (Ultralytics LLC, Washington D.C., USA). We trained the model using a dataset of 134 randomly selected images from multiple days of filming, including images from all cameras. We manually annotated each training image in Label Studio 1.10 (Heartex, San Francisco, USA) using the “Object Detection with Bounding Boxes” tool, with each of the 24 unique tags designated as a separate object class.

To train the model we resized each image to 640 × 640 pixels. Training took place over 300 epochs with a batch size of 8 and a learning rate of 0.01. To reduce false positive detections, we set a confidence threshold of 0.95, ensuring that only detections of >95% certainty were returned by the model. We also implemented YOLOv5’s built-in image augmentation protocols, which rotate, flip, and translate images in the training set, creating new variations of the original labelled training images. This process functionally increases the size of the training data set, reduces likelihood of overfitting and simulates different tag orientations (Zhang et al., 2024). Image augmentation makes the model more robust to possible variations in image orientation due to cockroach movement and enhances the model’s ability to accurately identify tags in images it had not previously seen.

We then evaluated the model by comparing its predictions against a manual inspection of 32 images that were not part of the training dataset, selected randomly from various cameras and filming days. These test images contained a total of 269 tagged cockroaches. The model correctly identified 131 individuals (48.7%), misidentified 11 individuals (4.1%), and did not detect 127 individuals (47.2%). We found that reducing the degree of certainty increased the number of correctly identified individuals, but also the number of misidentified individuals. Since misidentification could dramatically affect an individual’s social network position, while non-detections are less impactful given our very large sample size (Silk et al., 2015), we favoured a high degree of certainty. We applied the trained YOLO model to all collected images sequentially using a custom Python script that automatically processed images across all camera folders.

### Statistical analysis

We stored the lists of identified individuals per image as 192.csv files (four trials, three humidities, four days, four groups), ranging from 1378 lines to 3719 lines each (due to the varying number of successful detections). Using the R package asnipe (Farine, 2013) we converted each list into a group by individual matrix and then into an association matrix, where strengths of associations between individuals were set as simple ratio indexes (the number of images two individuals were seen together divided by the total number of times the two individuals were seen; Cairns & Schwager, 1987). We then used the packages Matrix (Bates et al., 2025), igraph (Csardi & Nepusz, 2006), and tnet (Opsahl et al., 2010; Opsahl & Panzarasa, 2009) to calculate the undirected individual-level social network measures of degree (the number of unique associations, irrespective of association strength), strength (sum of all association strengths; a weighted measure which therefore accounts for the strength of associations rather than present/absent), clustering coefficient (the strength of associations within all compete triads the individual is involved in versus those in incomplete triads; also weighted), and closeness (the inverse of average weighted distance to all other nodes; also weighted). Individuals not seen in a given day were assigned NAs for all social network traits (n = 49). Those seen but without any connections had a degree and strength of zero, and were assigned a closeness of zero, but were not assigned a clustering coefficient (we entered “NA”) as their relative strength within complete triads could not be determined. This resulted in 4543 measures of degree, strength, and closeness, and 4530 measures of clustering coefficient. We also calculated the network-level measures of density (the sum of all associations in the association matrix), connection variance (the variance of individual strengths), global clustering coefficient (the ratio of weights within a connected triad to those in unconnected triads across the entire network), assortment by degree (the correlation between degree scores of the two individuals connected by each edge in the network), and average distance (weighted pathlength) between nodes. We had 192 measures for each of these traits. We therefore had four individual-level and five network-level social network measures. Of these there were three pairs where the individual and network-level trait represent the same type of connectedness (strength and density, individual and global clustering coefficients, and closeness and average distance, although for the final pair high values have opposite interpretations – high values of closeness indicate well connected, while high values of average distance indicate poorly connected). Of the other measures we chose, degree’s direct analogue at the network level (number of edges in the network) would be redundant with density, while connection variance and assortment by degree do not have a direct analogue at the individual level.

Before analysis we removed 16 observations where nymphs had been accidentally tagged and placed in the groups instead of females (for a single week two boxes had one nymph and one box had two nymphs). We fitted all statistical models in glmmTMB v 1.1.10 (Brooks et al., 2017). For each of the four individual-level traits we fitted a univariate model with the trait as a response variable, humidity as a continuous fixed effect (rescaled to have 30% as -1, 40% as 0, and 50% as 1 so that a positive effect means higher trait values at higher humidities, testing the effect of humidity on trait means for Question 1), shelf and trial as categorical fixed effects with four levels each (as control variables), random intercepts for individual ID and group ID, random slopes with each of these with humidity, and random intercepts for date. The model for closeness would not converge with any of the random slopes, and the mode for degree would not converge with the random slopes for individuals, so we removed them and assume the variance they account for is negligible. For all traits we used a Gaussian error structure, and for all except clustering coefficient we used the default link function; for clustering coefficient we used a logit link function (as clustering coefficient is bounded between 0 and 1 this is the preferable transformation; Warton & Hui, 2011). For strength only we also needed to use the “BFGS” optimisation algorithm to aid convergence (as opposed to the default nonlinear optimizer “nlminb”). As we had no interactions we calculated p values for the fixed effects with an ANOVA with type two sum of squares in the package car (Fox & Weisberg, 2019). For the random effects we used code from Schielzeth and Nakagawa (2022) to calculate the conditional repeatability of individual IDs (r_I_) and group IDs (r_G_) and the relative importance of the random slopes at each level (r^2^_S,I_ and r^2^ _S,G_ for individual and group levels respectively, addressing Question 2), setting the mean and variance of the × variable to 0 and 0.667 respectively, and using -1 to 1 as the range to calculate repeatabilities.

For our analysis of the network-level traits, we fitted one univariate model for each of the five traits with an identical structure (including normal error distributions) to those fitted to the individual-level data, except that we did not include individual-level random intercepts or slopes. Density used the default link function, connection variance a log link and the BFGS optimiser, global clustering coefficient a logit link and the BFGS optimiser, assortment by degree the default link and no random slopes to achieve convergence, while average distance used the default link and the BFGS optimiser. We again followed Schielzeth and Nakagawa (2022) to calculate the conditional repeatability of group IDs (r_N_) and the relative importance of the random slopes (r^2^_S,N_; for Question 2). This then allowed us to qualitatively compare our results to the results for the individual-level traits by looking at the main effects of humidity, the ranking of repeatability scores, and the relative magnitude of intercepts compared to slopes, and so answer Question 3.

## Results

At the individual level, only closeness was affected by humidity, showing a slight increase at lower humidities (Fig. 2 & Table 1). The other three individual traits show no relationship with humidity (Table 1, Figs. S1-3), and likewise none of the five network-level traits were affected by humidity (Table 2, Figs S4-8).

**Table 1.** Statistical results for the effect of humidity on each of the four individual social network measures. Degrees of freedom are one in all cases.

| Trait | Estimate | Standard Error | Chi-squared | P value |
| --- | --- | --- | --- | --- |
| Strength | 0.037 | 0.063 | 0.349 | 0.554 |
| Degree | -0.271 | 0.356 | 0.579 | 0.447 |
| Clustering coefficient | -0.028 | 0.083 | 0.114 | 0.735 |
| Closeness | -0.024 | 0.010 | 5.777 | 0.016 |

**Table 2.** Statistical results for the effect of humidity on each of the five network-level social network measures. Degrees of freedom are one in all cases.

| Trait | Estimate | Standard Error | Chi-squared | P value |
| --- | --- | --- | --- | --- |
| Density | 0.883 | 1.384 | 0.407 | 0.523 |
| Connection variance | 0.087 | 0.089 | 0.952 | 0.329 |
| Global clustering coefficient | -0.038 | 0.052 | 0.547 | 0.460 |
| Degree assortativity | 0.008 | 0.011 | 0.536 | 0.464 |
| Mean distance | 0.346 | 0.293 | 1.399 | 0.237 |

**Figure 2.**
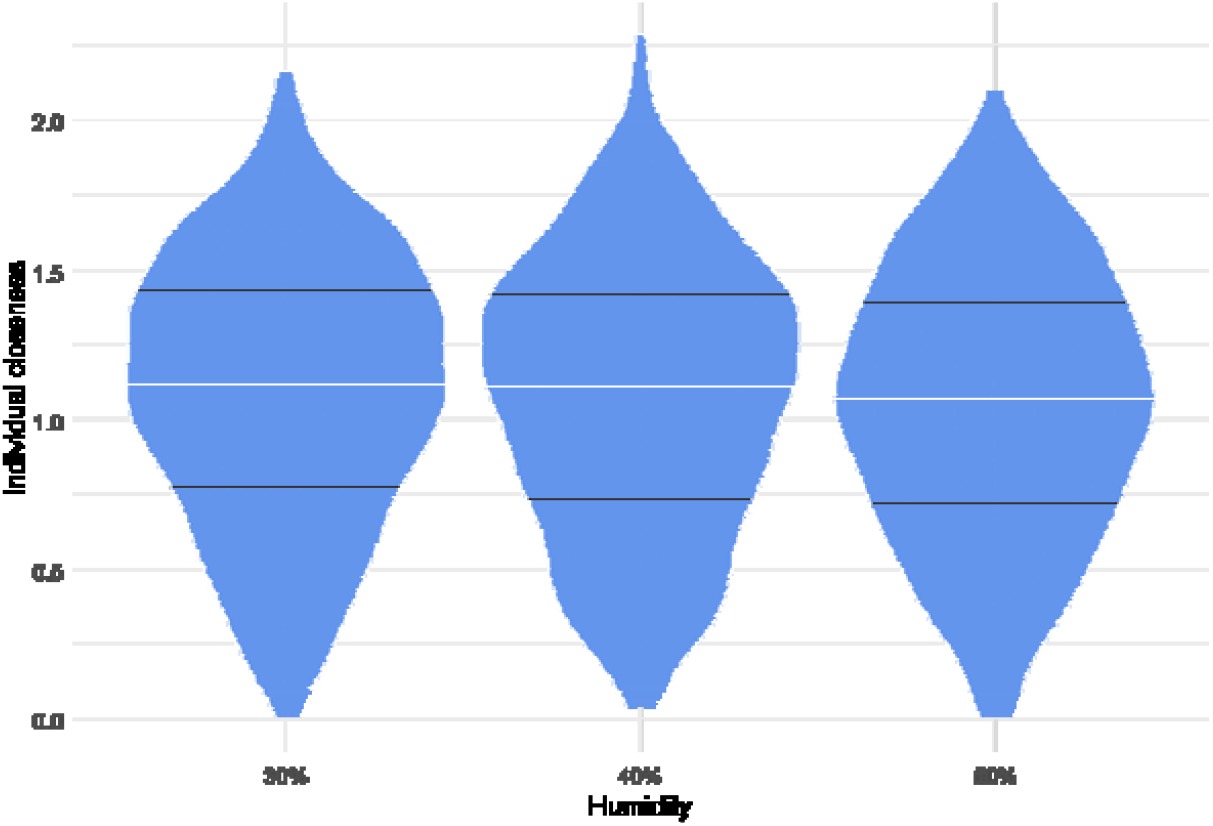
Individual closeness scores showed a slight decrease at the highest humidity. White lines indicate the median, while the black lines show the 25^th^ and 75^th^ percentiles.

For all four individual-level traits shelves differed in mean trait values (Table S1), with shelves 2 and 4 (the darker shelves) having a higher strength and lower degree, clustering coefficient, and closeness. Degree and closeness differed between trials too, with trial 1 having lower scores and trial 2 the highest (Table S2). For the network level traits, shelves 2 and 4 had higher connection variance, degree assortment, and average distance, but lower clustering coefficient, while degree did not differ among shelves (Table S3). All traits were highest in the first trial, except global clustering coefficient which did not vary among trials (Table S4).

For the individual-level analysis, individuals were somewhat repeatable in mean behaviours (r_I_ values were 0.115, 0.118, 0.387, and 0.155 for strength, degree, clustering coefficient, and closeness respectively), but their slopes are not consistently different (r^2^_S,I_ values were 0.088 for clustering coefficient and < 0.001 for three other traits, Fig. 3). In contrast, groups were less repeatable in their mean traits (r_G_ values were 0.017, 0.055, 0.263, and < 0.001 for strength, degree, clustering coefficient, and closeness respectively), but did show relatively larger consistent differences in slopes (r^2^_S,G_ values were 0.007, 0.049, 0.221, and 0 for strength, degree, clustering coefficient, and closeness respectively, Fig. 3).

**Figure 3.**
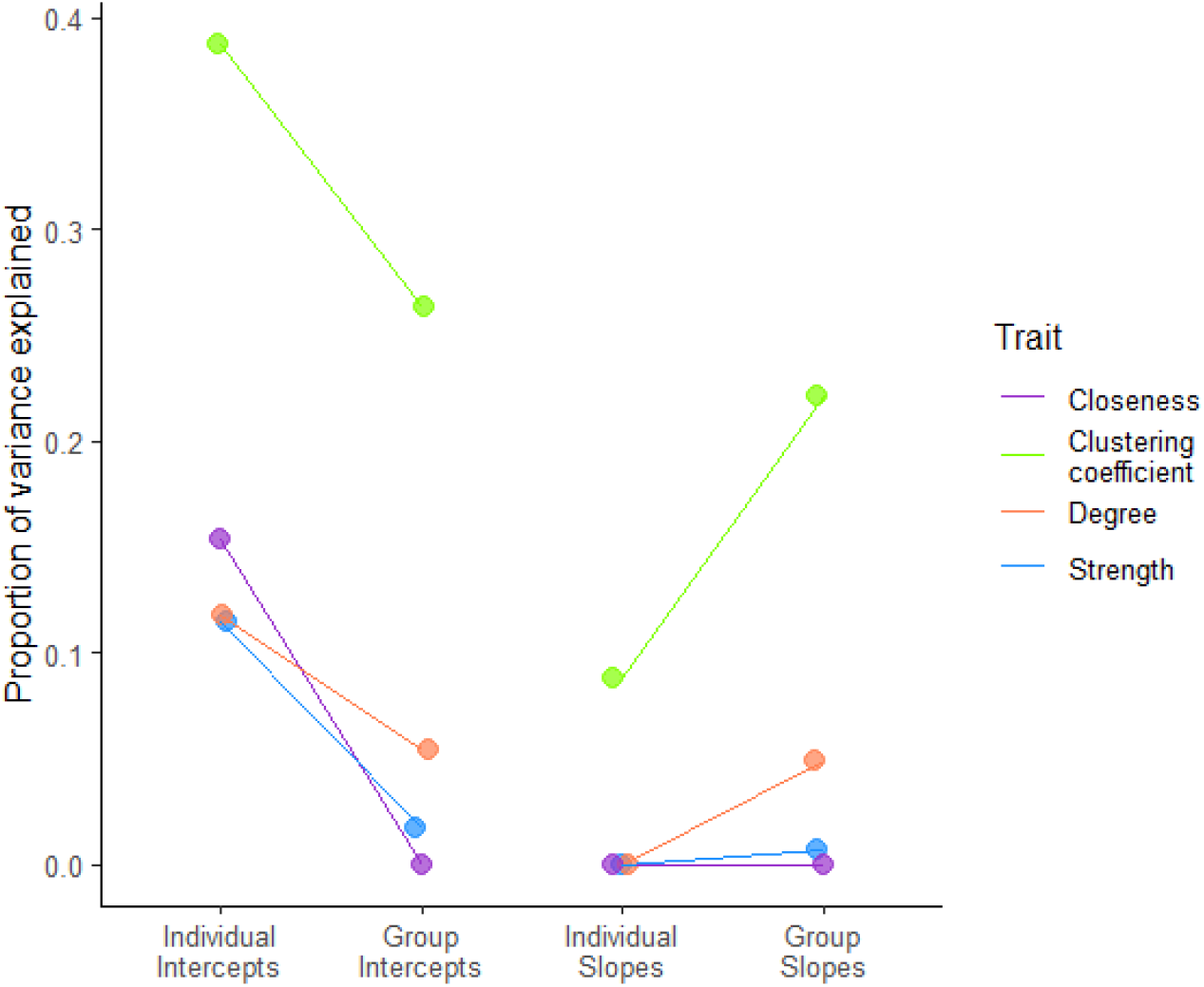
Estimates of the proportion of variance explained for both individual and group intercepts and slopes. Individual and group-level intercept repeatability estimates (r_I_ and r_G_) are connected by the same line (with different colours for each trait) to show the decline from individual level to group level, while the proportion of variance explained by individual and group-level slopes (r^2^_S,I_ and r^2^_S,G_) are connected by the same line to show the increase from individual level to group level.

In the whole network analysis, r_N_ values were 0.025, 0.022, 0.473, 0.011, and 0.060 for density, connection variance, global clustering coefficient, degree assortativity, and mean distance respectively. Values for r^2^_S,N_ were 0.010, 0.001, 0.464, 0.000, and 0.053 for density, connection variance, global clustering coefficient, assortativity, and mean distance respectively.

When comparing the patterns across the individual and network-level analyses, we see that groups have a relatively higher consistency in their plasticity, and a relatively lower consistency in their mean behaviours, than individuals do. The same trait (clustering coefficient) was the most repeatable at both the group-level of individual traits and for the entire-network traits, with closeness and mean distance the least repeatable (Fig. 4). Additionally, the darker shelves (2 and 4) had higher strength (individual level) and density (network-level), lower individual and global clustering coefficients (individual and network level respectively), and lower closeness (individual) and higher mean distance (network level) which are all compatible findings (Tables S1 & S3).

**Figure 4.**
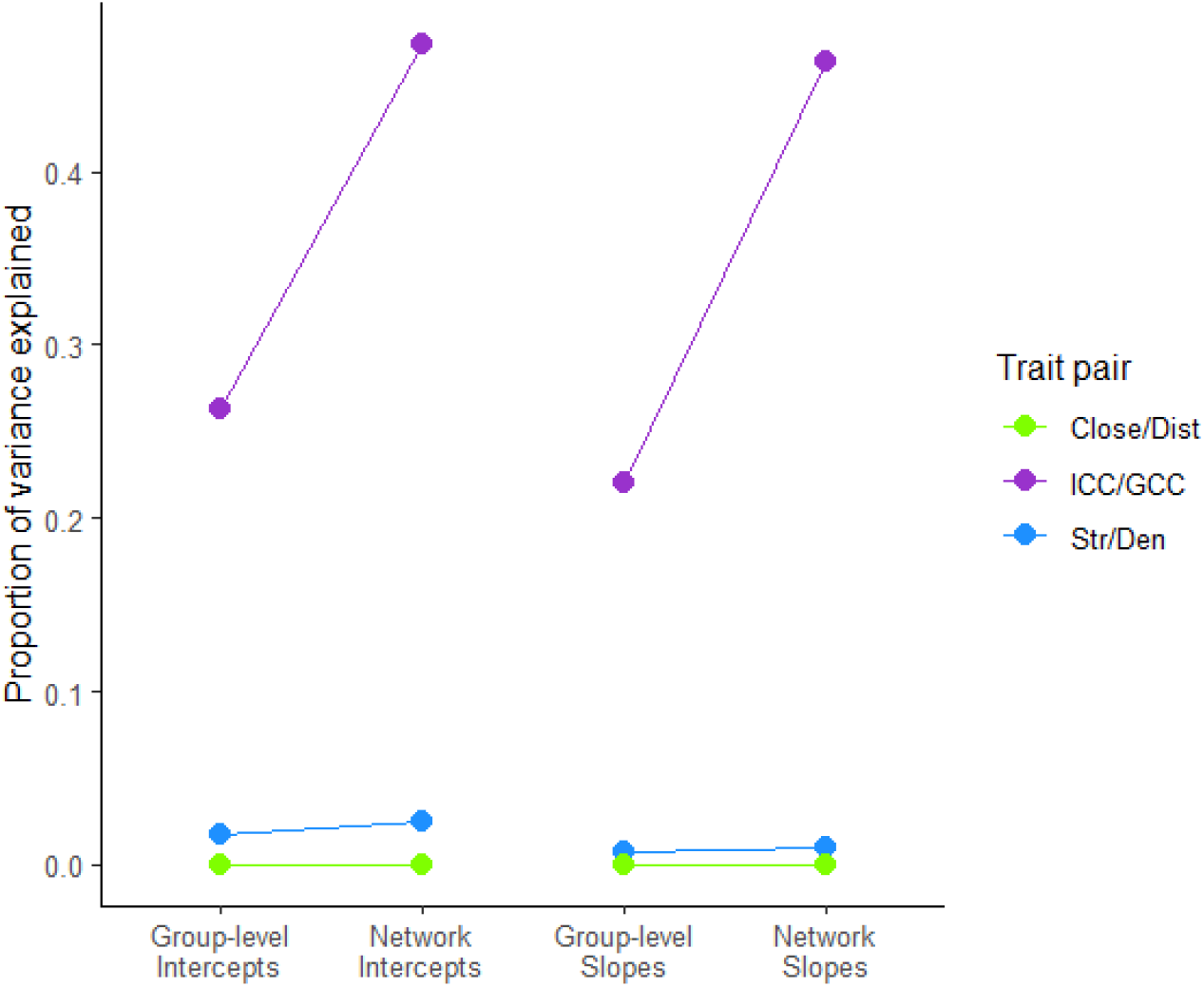
Estimates of the proportion of variance explained for the group-level variance components from the individual-level analysis (r_G_ [Group-level Intercepts] and r^2^_S,G_ [Group-level Slopes]) and the variance components from the network-level analysis (r_N_ [Network Intercepts] and r^2^_S,N_ [Network slopes]). Traits from the individual-level analysis (strength [Str], individual clustering coefficient [ICC], and closeness [Close] are connected by a solid line to their analogue at the network level (network density [Den], global clustering coefficient [GCC], and mean distance [Dist], respectively). This helps show that the rank order of traits does not change between the two analyses, indicating the approaches give comparable results.

## Discussion

We assessed change in individual and group-level social behaviour in cockroaches in response to a decrease in humidity but found very limited responses. Of the four individual-level social traits we measured, only closeness (connectedness to wider parts of the network) showed any clear change, with slightly increased closeness at lower humidities, but this effect was very small. Degree (number of unique connections), strength (overall gregariousness), and clustering coefficient (tendency of associates to also associate with each other) showed no clear change. We also saw no change in the network-level traits. We observed some consistency of individual mean behaviour, but not of individual plasticity, but in contrast we found limited group-level repeatability in means but more repeatability of plasticity.

Our finding of limited overall response to humidity suggests this cockroach species is robust to changes from 50 to 30% RH. Dambach and Goehlen (1999) found change in aggregative behaviour in response to humidity ranging from < 2 to >95% in first instar nymphs of the cockroach *Blatella germanica*, suggesting we may have seen a response if we had used a wider range of humidities. Since the current study, we have observed differences in burrowing behaviour between 20 and 80% RH in *B. dubia* (Profitt and Fisher, unpublished data), supporting the idea that this species is sensitive to humidity, but at a wider range than we tested here. Cockroaches in groups lose water at a slower rate than those isolated (Yoder & Grojean, 1997) and so by assaying ours in groups we may have given them the opportunity to buffer the 30% humidity sufficiently. At the highest humidity cockroaches had slightly lower closeness, which may reflect a reduced cost of being socially isolated in the most benign conditions (but note that both increased and decreased grouping can be expected in response to elevated environment stress; reviewed in: Fisher et al., 2021). However, this is a small effect made significant by our large sample size (see Fig. 2). If newer technologies such as those we used here become more common, and so sample sizes increase (through both larger experimental sample sizes and more efficient data extraction giving larger final datasets), we must maintain awareness of the difference between statistically significant and biologically significant results. Ogino *et al*. (2023) found environmental factors explained a considerable amount of variance (36-71%) in individual zebra finch (*Taeniopygia guttata*) social network traits, a magnitude of response much more likely to be biologically significant. We await further studies before we can start to identify general patterns e.g., responses only in certain taxa or due to particular environmental factors.

Alongside the limited plasticity, we found a modest degree of repeatability in individual social behaviours (ranging from 12 to 39% of the total variance). These are slightly higher than previous estimates of social behaviour in this population (8%; Fisher, 2023) and in line with mean estimates for invertebrates assayed in laboratories (24%; Bell et al., 2009). Cockroaches show consistency in individual means of various behaviours including boldness (Stanley et al., 2017), learning and memory (Arican et al., 2020), and thigmotaxis (Laurent Salazar et al., 2018), and so this is no surprise. What has been less well studied is consistency in individual plasticity. We found this was absent, meaning that sometimes an individual decreased their sociality in response to a drop in humidity, but sometimes they would increase. Individual variation in plasticity is expected to have a range of ecological and evolution consequences, such as modulating group responses to stimuli and setting the upper limit for the evolvability of plasticity (Dingemanse et al., 2010; Nussey et al., 2007). It is therefore interesting these were largely absent for social behaviour in *B. dubia*. If our cockroaches were truly not stressed by the range of humidities we exposed them to, as we suggest above, then the patterns we see could be explained by cockroaches not responding to humidity (no main effect of humidity), leaving individuals to differ due to underlying genetic or non-genetic but permanent factors like early life effects (repeatability of individual mean behaviour), alongside random fluctuations by individuals (limited consistency of individual slopes). If individuals genuinely do not differ in plasticity, then when exposed to more extreme RH values than we tested here they may all change in the same direction and to the same degree. This could lead to large shifts in group/population values, possibly leading to instability (Graham et al., 2006). The lack of consistency in plasticity in response to environmental stressors also suggests that plasticity itself is not heritable and so would not evolve regardless of any selective benefits to being more or less plastic.

In contrast to our findings for individual-level traits, group means showed limited repeatability, while group plasticity showed relatively more repeatability. Zebra finches also show less group-level repeatability in mean social network position than at the individual level (Ogino et al., 2023), but repeatability of plasticity was not quantified. More similar to our results, Gordon *et al*. (2023) found variation in collective plasticity in response to humidity in the red harvester ant (*Pogonomyrmex barbatus*). The variation in collective plasticity has been suggested to be key for allowing population persistence in the face of drought events (Gordon, 2013), but how such group-level variation can exist without individual-level variation in plasticity is less clear. In ants, collective day-night rhythms show different evolutionary patterns to individual day-night rhythms (Doering et al., 2024). This study suggests that, in these eusocial insects, individual and group-level behaviours can be decoupled, somewhat analogous to what we observed here.

In many systems of interacting entities, the system-level behaviour (“macrostate”) cannot be reduced to the behaviour of the individual entities (“microstates”), known as an “emergent phenomenon” or “weak emergence” (Bedau, 1997). This non-reducibility is what we found here, with no consistent individual plasticity yet consistent group-level plasticity. The most often proposed cause for emergent effects (across a range of meteorological, technical, and biological systems) is that the link between the microstate and microstate in the system features subtle nonlinear responses (such as a small change by one individual leading to larger changes in others) and/or feedback loops (e.g., one individual clustering less leads to other individuals clustering less) (Boeing, 2016; Fisher & Pruitt, 2019). This process is known as “deterministic chaos”, where deterministic, predictable rules give rise to unpredictable dynamics at larger scales (May, 1976; May & Oster, 1976). Therefore, it could be that there are very small (even imperceptible) differences among individual cockroaches which are then amplified by interaction dynamics to give a detectable signal (repeatability of plasticity) at the group level: some groups consistently increase their social grouping (for example) with humidity, while other decrease. For instance, if individuals initially differed by small amounts in willingness to join and leave groups, and they tended to copy the behaviour of others (a positive feedback loop, e.g. copying shelter choice: Lihoreau & Rivault, 2011), you might see larger changes in groups (as individuals keep joining or leaving aggregations over the trial) than are detectable at the individual level.

If the above explanation is true the individual-level differences would be too small to detect (hence no among-individual variation in plasticity) and at the population level there is no consistent change as groups change in different directions (hence no main effect of humidity), but at the group level there is consistency in change. If the change occurs continually that might also explain why there is limited group-level consistency in mean behaviour. Lyapunov exponents can be used to detect divergent trajectories in time series, which can indicate the presence of chaotic dynamics such as those we discuss here (Wolf et al., 1985), while agent based models may also be used to simulate emergent dynamics (Medina et al., 2021). The large datasets of many individuals interacting in many groups that we require to test these ideas of complexity and chaos are now within our reach thanks to machine vision approaches such as the one we use here. To assess the role of chaotic determinism in governing collective behavioural dynamics, future studies should specifically test the short and long-term predictability of group level behaviour, measure the trajectories of behaviour among groups to see if they diverge, and determine if “strange attractors” (areas of phenotypic space trajectories tend to converge around) exist for collective behaviours (Fisher & Pruitt, 2019; Hastings et al., 1993; see also: Ouellette, 2022).

We also found that patterns of consistency for means and plasticities of behaviour were similar between the group-level analysis of individual traits and the network-level analysis. Both sets of analyses also detected no change due to humidity, but the darker shelves resulted in lower individual and global clustering coefficient, and lower closeness and higher mean distance, results which all are directly comparable to one-another. These results indicate that cockroaches interacted in less well-connected groups in the dark, possibly as the cockroaches were not moving around to look for more shelter.

Alongside the results mentioned in the previous paragraph, we also found comparable patterns in variance between group-level variance in individual traits and network-level variance: closeness (individual) and distance (network) were the least repeatable, followed by strength (individual) and density (network), while clustering coefficient at both the individual and network level was the most repeatable. Researchers often either measure a group’s phenotype as the mean phenotype of individuals within the group or measure a property of the entire group that cannot be reduced to the level of the individuals. Here we show that these approaches should be comparable (at least for social network traits), aiding better comparison of studies moving forward. Alongside social networks, another type of often-quantified group-level trait are movement-based traits such degree of cohesion. It would be interesting to know if such group-level traits have analogues at the individual level that would allow a similar comparison as we conducted here.

Finally, we note that machine vision algorithms are continually improving. Improved algorithms may be able to collect more information from each image or process images faster. Additionally, new capabilities are being unlocked, such as marker-less individual recognition (Ferreira et al., 2020), the automatic detection of social interactions (Robie et al., 2017; Schofield et al., 2023), and image recognition in more complex environments such as in the wild is becoming more feasible (Wiltshire et al., 2023). The rate of data collection and the complexity of such data will only increase, and so it is crucial behavioural ecologists are trained to collect, analyse, and interpret these data suitably to allow important questions to be robustly addressed.

## Acknowledgements

We thank Keith Lockhart for his hard work in maintaining the cockroach stock population since 2021 and insightful suggestions for improving animal husbandry. This work was funded by NERC (grants NE/X013227/1 and NE/X018407/1).

