## Supplementary materials for "Multilevel-consistency of social behaviour in a cockroach"

### Supplementary Figures

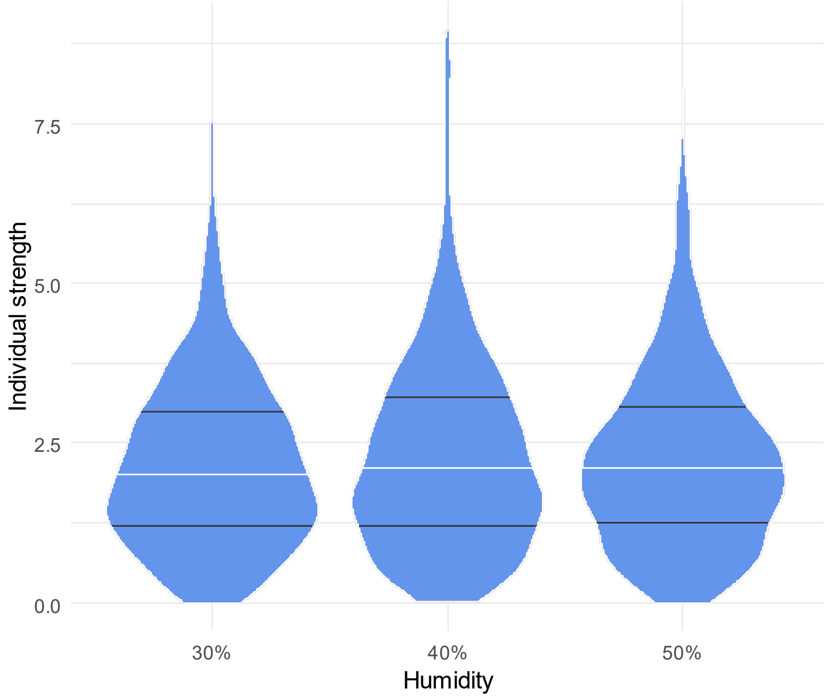


Figure S1. Individual strength scores showed no relationship with humidity. Width of each shape represents the frequency of scores, white lines indicate the median, and the black lines show the 25$^{th}$ and 75$^{th}$ percentiles.

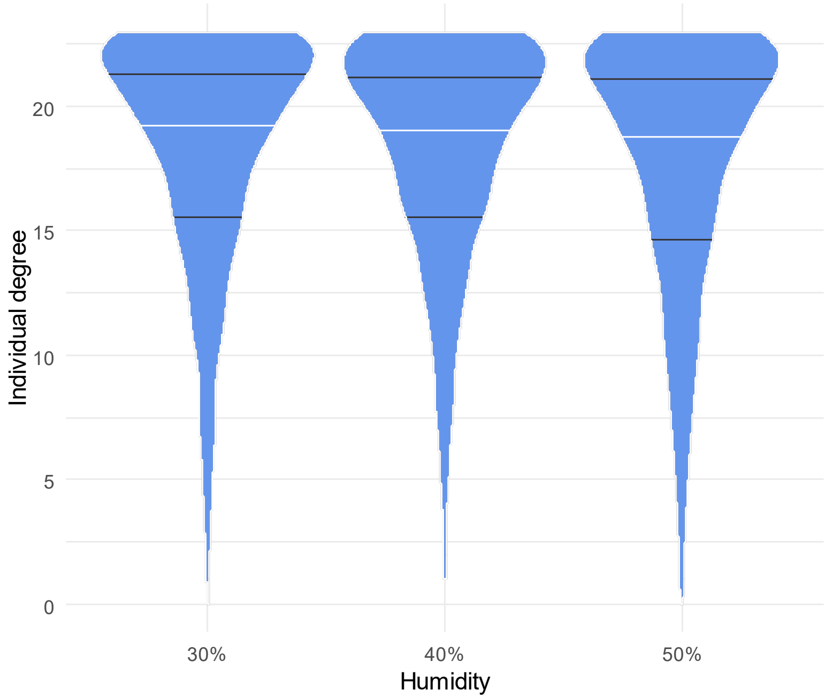


Figure S2. Individual degree scores showed no relationship with humidity. Width of each shape represents the frequency of scores, white lines indicate the median, and the black lines show the 25$^{th}$ and 75$^{th}$ percentiles.

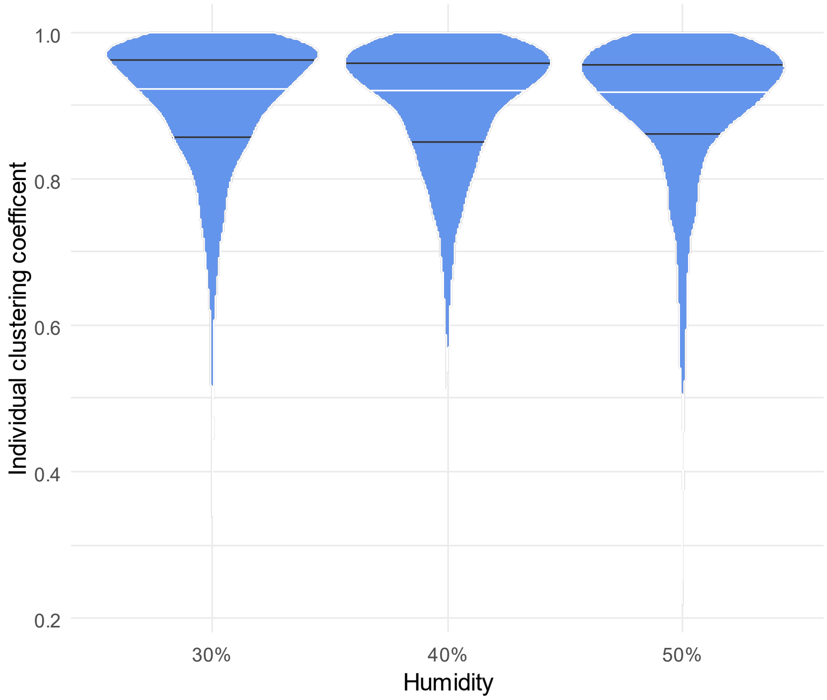


Figure S3. Individual clustering coefficient scores showed no relationship with humidity. Width of each shape represents the frequency of scores, white lines indicate the median, and the black lines show the 25$^{th}$ and 75$^{th}$ percentiles.

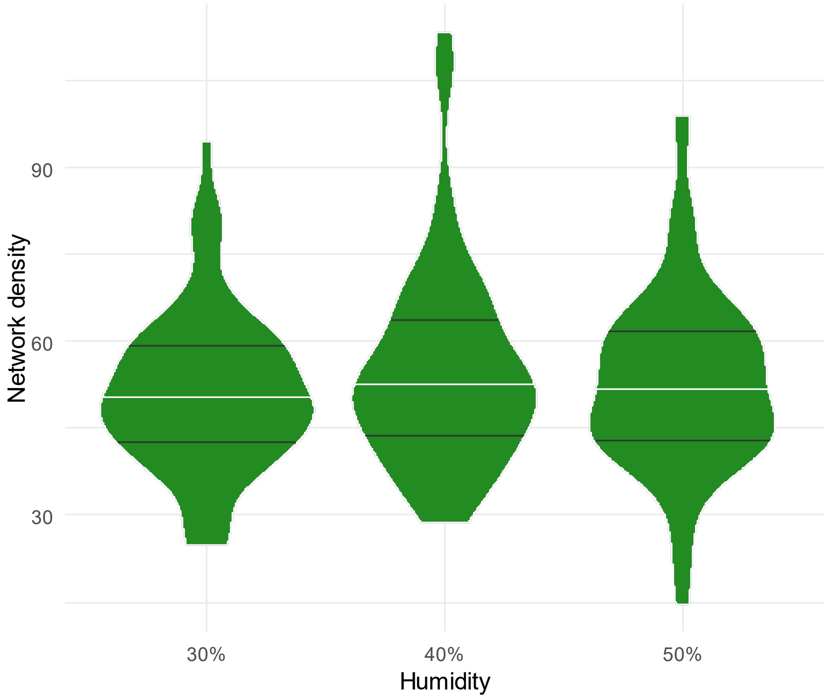


Figure S4. Network density scores showed no relationship with humidity. Width of the shape indicates frequency of scores, white lines indicate the median, and the black lines show the 25th and 75th percentiles.

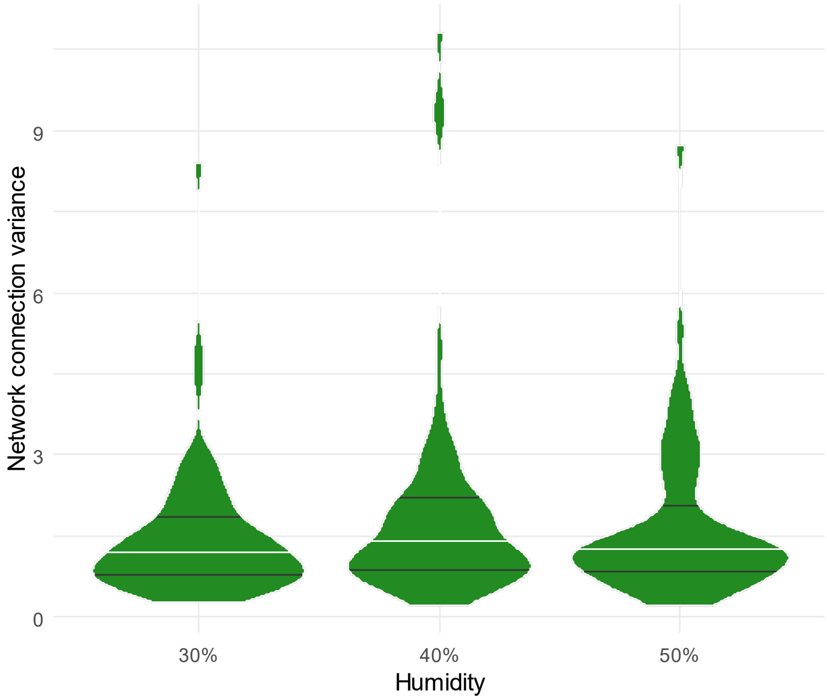


Figure S5. Network connection variance scores showed no relationship with humidity. Width of the shape indicates frequency of scores, white lines indicate the median, and the black lines show the 25th and 75th percentiles.

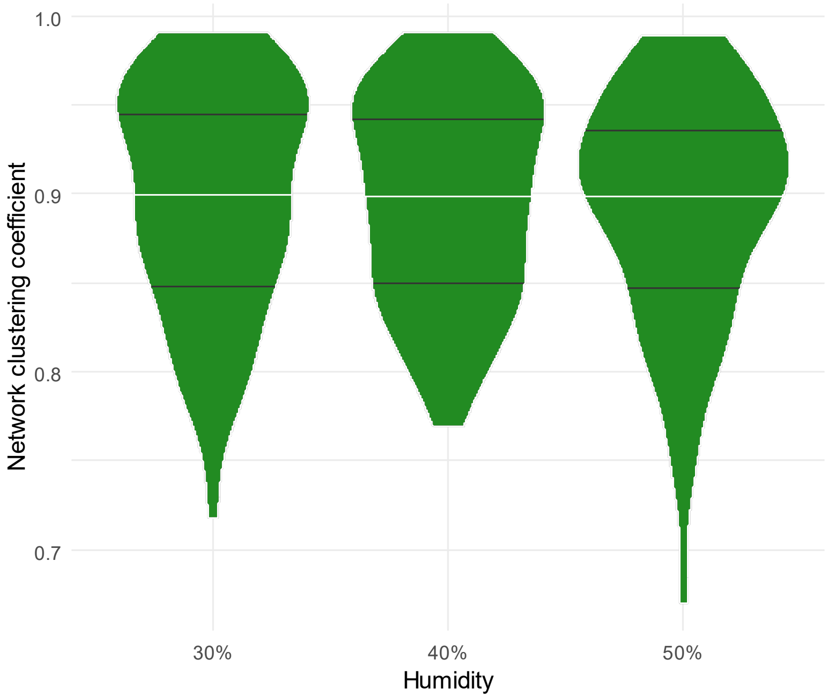


Figure S6. Network clustering coefficient scores showed no relationship with humidity. Width of the shape indicates frequency of scores, white lines indicate the median, and the black lines show the 25th and 75th percentiles.

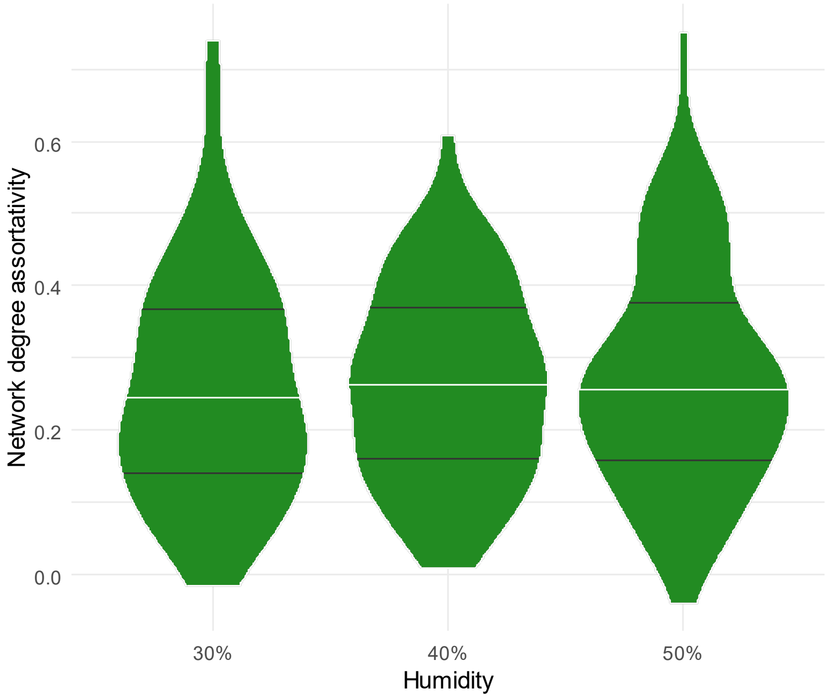


Figure S7. Network degree assortativity scores showed no relationship with humidity. Width of the shape indicates frequency of scores, white lines indicate the median, and the black lines show the 25th and 75th percentiles.

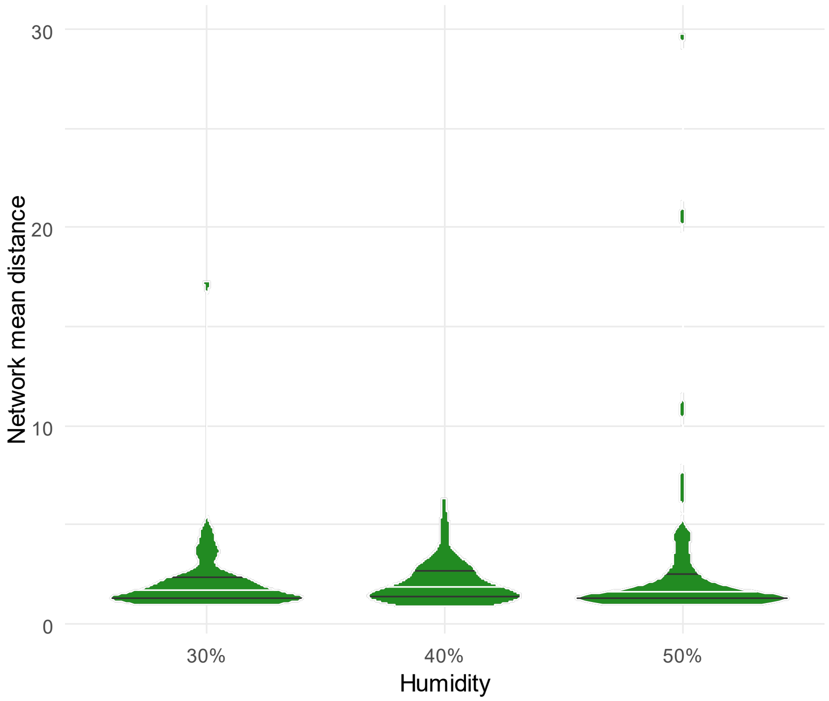


Figure S8. Network mean distance scores showed no relationship with humidity. Width of the shape indicates frequency of scores, white lines indicate the median, and the black lines show the 25$^{th}$ and 75$^{th}$ percentiles.

### Supplementary Tables

**Table S1**. Statistical results for the effect of shelf on each of the four individual social network measures. We set shelf 1 as the default and so the estimates indicate the difference between the other shelves and shelf 1. There is a single likelihood ratio test for each trait, testing the null hypothesis that the shelves have the same means. Degrees of freedom are three in all cases.

| Trait | Shelf | Estimate | Standard Error | Chi-square | P value |
|---|---|---|---|---|---|
| Strength | 2 | 0.179 | 0.054 | 20.855 | 0.000 |
| | 3 | -0.045 | 0.054 | | |
| | 4 | 0.101 | 0.054 | | |
| Degree | 2 | -3.170 | 0.156 | 1228.955 | 0.000 |
| | 3 | 0.385 | 0.155 | | |
| | 4 | -4.077 | 0.157 | | |
| Clustering coefficient | 2 | -0.808 | 0.043 | 793.680 | 0.000 |
| | 3 | 0.095 | 0.053 | | |
| | 4 | -0.860 | 0.043 | | |
| Closeness | 2 | -0.162 | 0.017 | 288.493 | 0.000 |
| | 3 | 0.021 | 0.017 | | |
| | 4 | -0.217 | 0.017 | | |

**Table S2**. Statistical results for the effect of trial on each of the four individual social network measures. We set trial 1 as the default and so the estimates indicate the difference between the other trials and trial 1. There is a single likelihood ratio test for each trait, testing the null hypothesis that the trials have the same means. Degrees of freedom are three in all cases.

| Trait | Trial | Estimate | Standard Error | Chi-squared | P value |
|---|---|---|---|---|---|
| Strength | 2 | -0.138 | 0.153 | 6.702 | 0.082 |
| | 3 | -0.213 | 0.153 | | |
| | 4 | -0.394 | 0.155 | | |
| Degree | 2 | 1.668 | 0.539 | 9.830 | 0.020 |
| | 3 | 1.058 | 0.540 | | |
| | 4 | 0.808 | 0.542 | | |
| Clustering coefficient | 2 | 0.283 | 0.137 | 5.290 | 0.152 |
| | 3 | 0.242 | 0.138 | | |
| | 4 | 0.107 | 0.137 | | |
| Closeness | 2 | 0.116 | 0.024 | 29.702 | 0.000 |
| | 3 | 0.083 | 0.024 | | |
| | 4 | 0.114 | 0.025 | | |

**Table S3**. Statistical results for the effect of shelf on each of the five network-level social network measures. We set shelf 1 as the default and so the estimates indicate the difference between the other shelves and shelf 1. There is a single likelihood ratio test for each trait, testing the null hypothesis that the shelves have the same means. Degrees of freedom are three in all cases.

| Trait | Shelf | Estimate | Standard Error | Chi-squared | P value |
|---|---|---|---|---|---|
| Density | 2 | 3.478 | 3.020 | 2.825 | 0.419 |
| | 3 | -1.318 | 3.020 | | |
| | 4 | 1.623 | 3.021 | | |
| Connection variance | 2 | 0.446 | 0.149 | 27.316 | 0.000 |
| | 3 | -0.460 | 0.233 | | |
| | 4 | 0.448 | 0.153 | | |
| Global clustering coefficient | 2 | -0.693 | 0.122 | 83.959 | 0.000 |
| | 3 | 0.117 | 0.156 | | |
| | 4 | -0.798 | 0.120 | | |
| Degree assortativity | 2 | 0.119 | 0.026 | 69.518 | 0.000 |
| | 3 | -0.015 | 0.026 | | |
| | 4 | 0.160 | 0.026 | | |
| Average distance | 2 | 1.394 | 0.521 | 24.916 | 0.000 |
| | 3 | -0.147 | 0.521 | | |
| | 4 | 2.024 | 0.522 | | |

**Table S4**. Statistical results for the effect of trial on each of the five network-level social network measures. We set trial 1 as the default and so the estimates indicate the difference between the other trials and trial 1. There is a single likelihood ratio test for each trait, testing the null hypothesis that the trials have the same means. Degrees of freedom are three in all cases.

| Trait | Trial | Estimate | Standard Error | Chi-squared | P value |
|---|---|---|---|---|---|
| Density | 2 | -2.801 | 3.385 | 10.315 | 0.016 |
| | 3 | -5.426 | 3.309 | | |
| | 4 | -10.252 | 3.369 | | |
| Connection variance | 2 | -0.760 | 0.231 | 16.626 | 0.001 |
| | 3 | -0.529 | 0.216 | | |
| | 4 | -0.772 | 0.233 | | |
| Global clustering coefficient | 2 | 0.209 | 0.119 | 4.904 | 0.179 |
| | 3 | 0.207 | 0.119 | | |
| | 4 | 0.050 | 0.113 | | |
| Degree assortativity | 2 | -0.098 | 0.027 | 14.134 | 0.003 |
| | 3 | -0.045 | 0.027 | | |
| | 4 | -0.070 | 0.027 | | |
| Mean distance | 2 | -1.889 | 0.555 | 14.785 | 0.002 |
| | 3 | -1.653 | 0.546 | | |
| | 4 | -1.680 | 0.548 | | |
